# Determining the toxicity and potential for environmental transport of pyridine using the brown crab *Cancer pagurus* (L.)

**DOI:** 10.1101/2022.11.17.516169

**Authors:** Chloe L. Eastabrook, Miguel Morales Maqueda, Charlotte Vagg, Joyce Idomeh, Taskeen A. Nasif-Whitestone, Poppy Lawrence, Agnieszka K. Bronowska, John H. Bothwell, Brett J. Sallach, Joe Redfern, Gary S. Caldwell

## Abstract

A series of mass mortalities (wash-ups) of marine life were documented along England’s north east coastline with peaks in September and October 2021, coincident with a programme of intensified maintenance dredging of the River Tees. Decapod crustaceans were the worst affected fauna, with brown crab (*Cancer pagurus*), European lobster (*Homarus gammarus*, L.), green shore crab (*Carcinus maenas*, L.) and velvet swimming crab (*Necora puber*, L.) populations severely affected. Moribund animals presented with twitching behaviours and paralysis. A potential release of the industrial pollutant pyridine was forwarded as one explanation; however, toxicology data for pyridine in decapods is lacking. In this study, we address this knowledge gap by executing a programme of immersion exposure experiments (pyridine at 2 - 100 mg L^-1^) using *C. pagurus*, measuring toxicity effects at the individual (survival) and cellular levels (cellular, mitochondrial, and lipid peroxidation reactive oxygen species (ROS) formation in the gills, hepatopancreas and claw muscle). Highest mortality rates were seen after 72 hours of exposure, returning an LC_50_ value of 2.75 mg L^-1^. Exposed crabs presented with patterns of convulsions, limb twitching, paralysis, and death. Crabs exposed to the lowest pyridine dose (2 mg L^-1^) were noticeably more docile than controls. Concentration was a significant factor influencing mitochondrial ROS formation at low concentrations, with tissue type, time, and their interaction all significant at 100 mg L^-1^. Computer simulations were used to model the transport of any pyridine released from the dredging work, demonstrating the potential for a pyridine plume to extend from Seaham to the north of the Tees to Whitby and Robin Hood’s Bay to the south. This range corresponds well with the reported wash-ups and subsequent declines in catch rates.

## 2 Introduction

England’s North East and North Yorkshire coastlines experienced major marine mass mortality events during September and October 2021, affecting limpets, octopuses, mussels, razor clams, seabirds, seals, and porpoises (Das, 2022). However, the large decapod crustaceans were disproportionately affected, including commercially important species such as brown crabs (*Cancer pagurus*, L.), European lobsters (*Homarus gammarus*, L.); with other ecologically important crustaceans heavily impacted most notably green shore crabs (*Carcinus maenas*, L.) and velvet swimming crabs (*Necora puber*, L.). These high incidences of crustacean deaths have since become synonymous with these events (Isaac, 2022). Mortalities were evidenced by a series of ‘wash-ups’ of dead and moribund crustaceans along local beaches, and the region’s fishing industry reported dramatic and sudden declines in shellfish landings (NEIFCA, 2022). Aside from wash-ups, crabs and lobsters were also reported to be found dead and dying inside pots placed in inshore waters. Those shellfish that did survive capture were commonly reported to have died *en masse* in the holding tanks of established buyers within the local seafood supply chain (NEIFCA, 2022).

Moribund crabs presented specific behavioural traits, notably lying supine and exhibiting pronounced twitching movements of the claws and walking legs. These behaviours were widely reported by representatives of the local fishing industry, recreational scuba divers, and a diverse spectrum of coastal stakeholders. The epicentre of the observed mortalities were centred on the Hartlepool, Teesmouth and Redcar areas of the Cleveland coast; however, impacts were observed by fishing fleets along almost 50 km of the north east coastline, stretching from Scarborough in North Yorkshire to the south and Seaham in County Durham to the north (NEIFCA, 2022). Mortalities occurred in two phases: first, an acute phase that accounted for the initial high numbers of deaths, and second, a more chronic albeit still relatively fast-acting phase that affected surviving crustaceans both at sea and after having been caught and sold into the seafood supply chain. This more chronic phase (perhaps resulting from secondary toxicity or because of an overall reduction of fitness) likely explained the series of wash-ups reported in late October and into November 2021.

Initial investigations conducted by agencies of the UK Government’s Department of Environment, Food & Rural Affairs (DEFRA) concluded that a satellite-visible algal bloom seen off Teesside in Sep 2021 was likely to have been responsible (DEFRA, 2022), either through the direct production of biotoxins or as a result of deoxygenation following bloom collapse. However, no direct monitoring of the bloom was carried out and evidence for its status as a harmful algal bloom remains circumstantial (Deere-Jones, 2022). An alternative explanation is that the mortalities were triggered in response to a period of increased maintenance dredging at the mouth of the River Tees shipping channel that may have released industrial pollutants sequestered in the sediment. The onset of the mass mortalities was coincident with an intensive 10-day dredging campaign by the trailing suction hopper dredger UKD Orca, in response to slippage of sediment into the shipping channel (EFRA, 2022). The UKD Orca was in continuous operation, completing 52 dredging events and transporting around 150,000 tonnes of sediment and sediment-associated water to spoil dumping grounds.

Further data contained in the Defra reports indicated that tissues of the dead crabs contained elevated levels of pyridine at up 439.6 mg kg^-1^, versus 5 mg kg^-1^ for reference crab samples collected from Penzance in Cornwall, southwest England, albeit using a non-validated detection method (DEFRA, 2022). A report commissioned by the local fishing industry (Deere-Jones, 2022) considered whether pyridine could have been a causative agent given pyridine’s prevalent use in, and discharge from, historic and contemporary Teesside industries.

Pyridine, a heterocyclic organic compound, is a widely used industrial solvent and precursor chemical, contributing to the manufacture of several potent biocides (Onduka et al., 2013;El-Dean et al., 2019). It is also a high-volume by-product of some large industries including oil refining and the coking process used for steelmaking (Lee et al., 1991;Bai et al., 2011;Elkasabi et al., 2014), both of which are well represented in the industrial heritage of the Tees area (Warren, 1969;Molle and Wever, 1984). The available toxicology data for pyridine against aquatic organisms is, however, limited: examples include 96-hour LC_50_ values of 73.6, 26 and 4.6 mg L^-1^ for fathead minnow (*Pimephales promelas*), European carp (*Cyprinus carpio*), and rainbow trout (*Oncorhynchus mykiss*) (ThermoFisher, 2009), respectively.

Given the paucity of toxicological data for pyridine against aquatic organisms (Vaal et al., 1997), and the absence of pyridine-specific toxicological data for decapod crustaceans, the current work aimed to determine whether pyridine is toxic to *C. pagurus* and, if toxic, to characterise the nature of the toxicity. Further, on the assumption that pyridine could be liberated from dredged sediment, computational approaches were used to simulate the rate and geographical spread of pyridine transport along a coastline. We intend our work to inform debate around the probable causes of the mass mortality events and provide an evidence base for future monitoring, management, and mitigation strategies targeted at the Tees estuary and other marine ecosystems worldwide.

## 3 Methods

### 3.1 Crab husbandry

*Cancer pagurus* were collected by commercial potting from waters around Coquet Island, Co. Northumberland and landed at the Port of Blyth, north east England, during the months of June, July, September and October 2022. Upon landing, the crabs were immediately transported to the laboratory in insulated boxes where they were acclimated in a series of individual tanks containing 70 L of aerated artificial seawater (ASW) (pH 8.1 ± 0.2, salinity 35‰) with not more than seven crabs stocked per tank. The initial temperature was set at 11 °C and was incrementally increased by 1-2 °C every 2 days until tanks reached 16 °C. The light regime (16:8 hours light: dark) was controlled using low level fluorescent tubes. The crabs were acclimatised to these conditions for 7 days, during which they were twice fed to satiety using frozen squid. Uneaten squid was removed after 2 hours followed by a 50% water change. The crabs were regularly monitored for survival and signs of stress prior to the start of the experiments.

### 3.2 Pyridine exposure

The exposure tanks (8.9 L polypropylene) were filled with 4 L of artificial seawater. All pyridine experiments were conducted in chemical fume hoods with the temperature maintained at 16 ± 2°C using free-standing air-conditioning units. On the day of each experiment the tanks were placed inside the fume hoods and allowed to equilibrate to 16 °C. Once the exposure tanks were at the correct temperature, pyridine (99.8% purity, CAS-no: 110-86-1, Merck, UK) was added at concentrations between 2 - 100 mg L^-1^. After thorough mixing, a crab was added to each tank and a polypropylene lid was fitted. Each lid had two predrilled holes for an air line and air release. Tanks were aerated throughout the experiment using aquarium pumps fitted with air stones. There were no water changes and the crabs remained unfed.

### 3.3 Pyridine survival assay

Following an initial range finding experiment, a concentration response study was conducted to determine the No Observed Effect Concentration (NOEC), Low Observed Effect Concentration (LOEC) and the Lethal Concentration that results in 50% mortality (LC50) following pyridine exposure over 72 hours. Pyridine concentrations were set at 2, 5, 10, 20, 50 and 100 mg L^-1^. Control crabs were exposed to ASW only. Triplicates were used for each concentration. Survival was monitored at 24, 48 and 72 h and behavioural observations were noted. Survival was scored as 1 for alive and 0 for dead at each time point. Survival data were transformed and normalised to be expressed as percentages and plotted as a concentration response curve. Water and tissue samples were collected for subsequent pyridine analysis. All surviving crabs were humanely sacrificed by spiking at the end of the experiment after being placed on ice for 20 minutes.

### 3.4 Reactive oxygen species (ROS) assay

A separate experiment was conducted to determine ROS production. Crabs (n= 3) were exposed for 6 hours and 24 hours to four pyridine concentrations (2, 5, 20 and 100 mg L^-1^) alongside a control. Survival at each concentration was noted. Each crab was removed from its tank, placed on ice for 20 minutes to slow activity (Serrano and Henry, 2008), and humanely sacrificed by spiking. Each crab was sexed, the carapace width measured and whole crab weight was recorded prior to dissection. The gills and claw muscle were extracted and individually weighed for each time point before being placed into separate 50 mL falcon tubes containing 0.1M Tris HCL buffer at a 1:1 w/v (pH 7.4). Additional samples from the hepatopancreas were taken from the 6-hour exposure at 100 mg L^-1^. Each sample was homogenised using a benchtop homogeniser (Vevor high speed homogeniser FSH-2A, small blade) and centrifuged for 20 minutes at 3,000 *g*. The supernatant was collected and stored in Eppendorf tubes to be frozen at −20 °C until further analysis. Haemolymph samples were taken, but rapidly coagulated and were consequently discarded.

On the day of analysis, the tissue supernatant samples were defrosted on ice and 180 μL of each sample were added into a clear bottom 96-well microplate (Corning Costar). Cellular ROS (cROS), mitochondrial ROS (mROS) and lipid peroxidation ROS (LPO ROS) were determined *in situ* using the probe 2’7’-dichlorodihydrofluorescein diacetate (H2DCFDA, Invitrogen, Molecular Probes Inc., Eugene, OR, USA), dihydrorhodamine 123 (DHR123, Invitrogen, Molecular Probes Inc., Eugene, OR, USA) and BODIPY™ 581/591 C_11_ (Invitrogen, Molecular Probes Inc., Eugene, OR, USA), respectively. Stock solutions of H2DCFDA, DHR123 (20 mM) and BODIPY C_11_ were diluted to a working solution of 2 mM in MilliQ water in darkness, of which 20 μL was added to each well, with a final concentration of 200 μM. The microplate with cROS and mROS samples was analysed for ROS formation using a FLUOstar Optima fluorescence plate reader (BMG Labtech, Ortenberg, Germany) with an emission wavelength of 535 nm and excitation wavelength of 485 nm. The LPO ROS samples were measured at emission 620 nm and excitation 560 nm using a TECAN Spark 20m fluorescence microplate reader (TECAN, Männedorf, Switzerland). All ROS data were measured in relative fluorescent units (RFU) and normalised to the mass of tissue collected, and further normalised to the control samples to be presented as fold changes.

### 3.5 Computational modelling

The NEMO (Nucleus for European Modelling of the Ocean) modelling framework (Madec and NEMO-System-Team) was used to simulate the dispersion of a numerical tracer representing pyridine along the eastern coast of Great Britain. The NEMO framework includes a set of state-of-the-art routines for the simulation of ocean mean flow dynamics, the parameterisation of turbulence and the transport of active (temperature and salinity) and passive tracers (such as pyridine).

The model domain covers the spherical rectangle (48°N,3°W)–(58°N,3°E), as depicted in Figure S1. The integration grid within the domain is identical to the grid of the Atlantic Margin Model 1.5 km, AMM15 (Graham et al., 2018), for that area. The horizontal resolution is approximately 1.5 km and there are 33 levels in the vertical, 17 of which are in the upper 200 m of the water column.

Ocean dynamics in the domain are driven in the following way. Hourly surface currents and daily, three-dimensional temperatures and salinities in the domain were extracted from the AMM15 analysis simulation for the period 24^th^ September 2021 at 1200 to 1^st^ of November 2021 at 1200. The model AMM15 is used by the Met Office to forecast operational sea states in the Northwest European Shelf and represents our best estimates of ocean flows and properties in the area. The data were downloaded from the Copernicus Marine Service (NORTHWESTSHELF_ANALYSIS_FORECAST_PHY_004_013 product). These data were assimilated in NEMO by restoring the relevant variables in the model to the corresponding AMM15 fields with relaxation timescales of 900 seconds (one quarter of an hour) for the surface currents and 86400 seconds (one day) for temperatures and salinities. This approach ensures that circulation in the model at all depths very closely follows that of the original AMM15 simulation, including the cycle of the 11 dominant tidal constituents.

A numerical passive tracer, nominally pyridine, was released in the model grid boxes nearest to the seafloor and closest to the areas of dredging (54.66°N,1.12°W, 17.5 m) and disposal (54.70°N, 1.03°W, 35 m). The rationale for this approach is that any pyridine released by dredging, being initially sorbed in the disturbed sediments, will probably be predominantly encountered near the seabed, where the sediment plume is expected to concentrate. Ten thousand litres of tracer were released at each of these two sites uniformly in time between 26-09-2021 at 1200 and 03-10-2021 at 1200. The chosen period of release coincides with the time during which UKD Orca was operating in the area https://voyage.vesselfinder.com/fc1a3a2314b1fa47175d58b9d77165fe). The amount of substance that was potentially released by dredging is not known, and so the quoted 10,000 L volume is largely indicative but follows that used by the Centre for Environment, Fisheries and Aquaculture Science (Cefas) in their original model of potential pyridine transport (DEFRA, 2022). The dynamics of tracer dispersion in the model are such that the relative distribution of tracer is not affected by the quantity of tracer released, because the ratio of the rate of change of tracer concentration at a given point and moment in time to the rate of tracer released does not depend on the rate of release if the latter remains uniform in time. After release, the tracer is advected by the simulated mean flow and turbulently diffused both horizontally and vertically. Vertical turbulence is parameterised according to Blanke and Delecluse (1993). Given that the release of pyridine occurs at the bottom of the water column and that the potential for pyridine volatilisation is small (ATSDR, 2011), pyridine air-sea exchanges were disregarded in this simulation. However, degradation/oxidation of pyridine was considered by postulating a local exponential decay of concentrations with an e-folding time of 3.47 days (ATSDR, 2011). This means that 90% of the released pyridine will have decayed after 8 days.

### 3.6 Statistical analysis

All statistical analyses were conducted using GraphPad Prism 9 (GraphPad Software, San Diego, USA). Using a simple logistic regression, survival data from 24 and 72 hours were plotted and the slope tested for non-zero significance. Alongside this, a non-linear regression was used to calculate the LC_x_ values, NOEC and LOEC values after 24 and 72 hours. All data were checked for normality using the Shapiro-Wilk normality test and analysed using a two-way analysis of variance (2W ANOVA), with Post-hoc Tukey test to determine significant differences between concentrations. ROS data were normalised to tissue mass and control RFU values, and then expressed as fold change data.

## 4 Results

### 4.1 Survival and behaviour

Survival remained high following 24 hours exposure at pyridine concentrations below 20 mg L^-1^; however, all crabs were dead after 24 hours at 50 and 100 mg L^-1^. There was no change in mortality between 24 and 48 hours, therefore only the 24-hour data are presented (Figure 1A). Further mortalities were recorded after 72 hours, with an LC_50_ value of 2.75 mg L^-1^ (Figure 1B; Table 1). Both time and concentration were significant factors affecting survival, but the interaction between time and concentration was not significant (Table 2). Both the 24- and 72-hour slopes were not significantly non-zero (P = 0.068 and P = 0.122 respectively).

**Figure 1.**
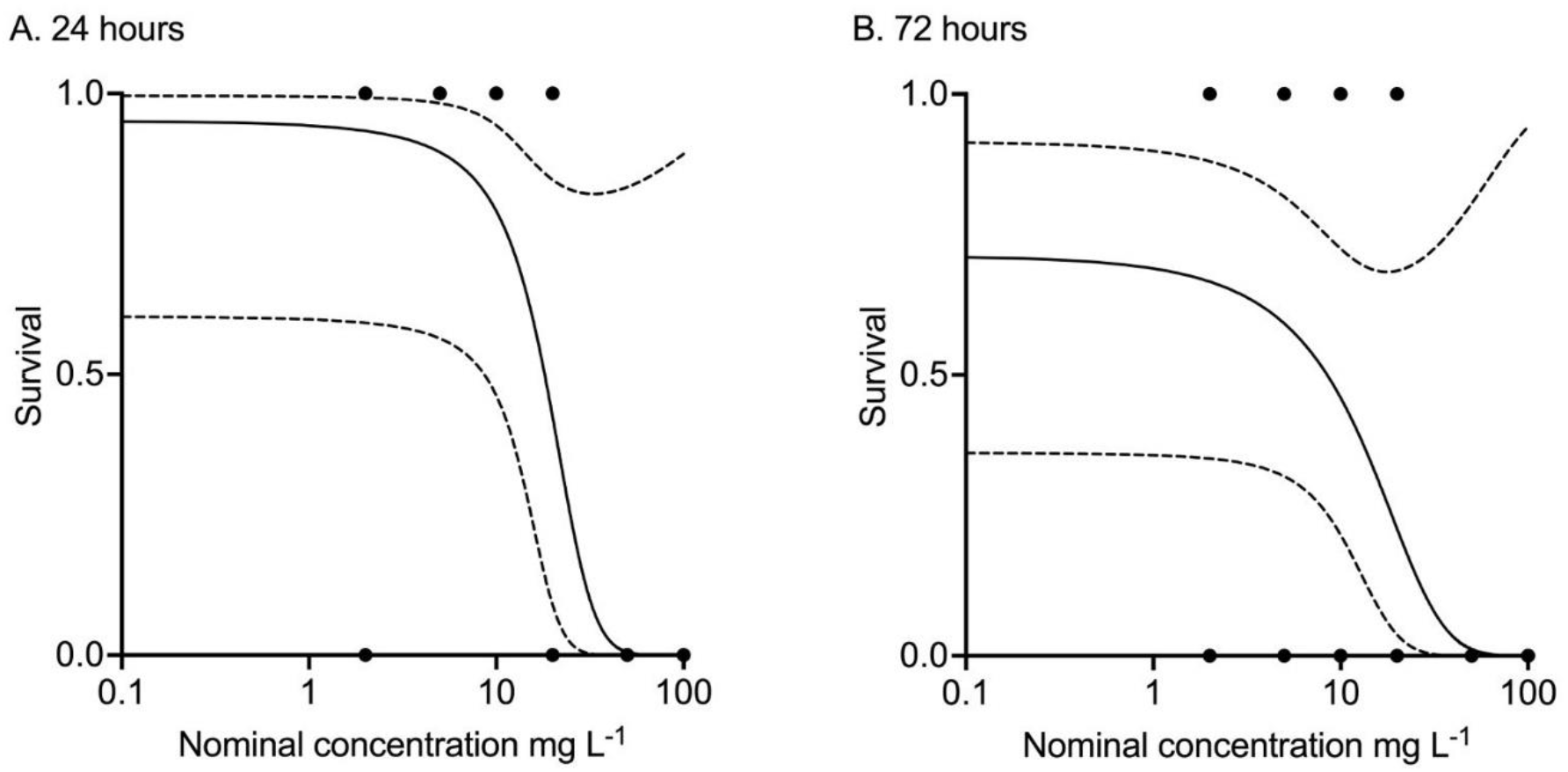
Concentration response curve for *Cancer pagurus* survival after (A) 24 hours (R^2^ = 0.65) and (B) 72 hours (R^2^ = 0.33) of exposure to pyridine (n = 3). The solid black line represents the logistic regression and the two dashed lines represent the confidence limits at 95%.

**Table 1.** Concentration response curve statistics for survival following 24 and 72 hours of pyridine exposure, presenting the lethal concentration that results in 10, 20 and 50% mortality (LC_10_, LC_20_ and LC_50_), No observed effect concentration (NOEC) and low observed effect concentration (LOEC) values. Values obtained from the non-linear regression (not presented).

| Time (hours) | LC<sub>10</sub> mg L<sup>-1</sup> | LC<sub>20</sub> mg L<sup>-1</sup> | LC<sub>50</sub> mg L<sup>-1</sup> | NOEC mg L<sup>-1</sup> | LOEC mg L<sup>-1</sup> | R<sup>2</sup> |
| --- | --- | --- | --- | --- | --- | --- |
| 24 | 17.38 | 18.11 | 19.44 | 20 | 50 | 0.69 |
| 72 | 0.066 | 0.26 | 2.75 | 20 | 50 | 0.37 |

**Table 2.** Two-way ANOVA statistics. * Denotes significant P values (P < 0.05). mROS = mitochondria reactive oxygen species, LPO = lipid peroxidation, conc. = concentration.

| Endpoint | Source | df | Sum of squares | Mean square | F | P |
| --- | --- | --- | --- | --- | --- | --- |
| Survival | Time | 1 | 0.3810 | 0.3810 | 5.333 | 0.0367* |
|  | Conc. | 6 | 5.4760 | 0.9127 | 4.259 | 0.0120* |
|  | Interaction | 6 | 0.6190 | 0.1032 | 1.444 | 0.2664 |
| mROS<br>(2 - 20 mg L<sup>-1</sup>) | Tissue | 1 | 0.2009 | 0.2009 | 0.888 | 0.3735 |
|  | Conc. | 3 | 0.9982 | 0.3327 | 4.860 | 0.0328* |
|  | Interaction | 3 | 0.2021 | 0.0674 | 0.298 | 0.8261 |
| mROS<br>(100 mg L<sup>-1</sup>) | Tissue | 2 | 284.5 | 142.2 | 6.506 | 0.0314* |
|  | Time | 2 | 296.4 | 148.2 | 8.612 | 0.0255* |
|  | Interaction | 4 | 503.7 | 125.9 | 7.317 | 0.0032* |
| LPO ROS<br>(100 mg L<sup>-1</sup>) | Tissue | 2 | 3.688 | 1.844 | 5.261 | 0.0479* |
|  | Time | 2 | 16.03 | 8.016 | 27.25 | 0.0002* |
|  | Interaction | 4 | 8.797 | 2.199 | 7.473 | 0.0029* |

A broadly concentration-dependent gradation of behaviours was observed. Crabs exposed to 100 and 50 mg L^-1^ presented twitching of the walking legs and in more extreme examples the crabs expressed convulsive-type responses. Onset of these motor responses was rapid at 100 mg L^-1^ (typically after 2-5 minutes of exposure). Individuals that presented violent convulsions performed repeated somersaults in their tanks. The convulsion phase was short-lived in the 100 mg L^-1^ treatment, lasting approximately 5-10 minutes. The crabs that had exhibited the somersault behaviours tended to come to rest upside-down and were unable to right themselves, presumably through weakness. These crabs then entered a ‘paralysis phase’ during which they lost the capacity to move their limbs over the next 20-30 minutes. At 100 mg L^-1^, all crabs were paralysed within 1 hour. The crabs were still alive as evidenced by the continued operation of their mouthparts. In all cases the crabs had died after 6 hours of exposure as evidenced by the cessation of any respiratory activity. Crabs exposed to 50 mg L^-1^ did not exhibit such extreme behaviours, but rapidly twitched their legs in a seemingly uncoordinated manner. Further, crabs exposed to 100 and 50 mg L^-1^ also tended to autotomise their claws and legs. This was spontaneous and not in response to any mechanical stimulation. Some loss of limbs was noted at lower pyridine concentrations but was not as prevalent as for the 100 and 50 mg L^-1^ treatments.

Crabs exposed to lower pyridine doses exhibited gradients of these behaviours. For example, the sole surviving crab at 20 mg L^-1^ was partially paralysed at the 24 hour observation, specifically the first two pairs of pereiopods and the chelae. This caused the crab to pitch forward, coming to rest on the fronto-orbital border of the carapace. The third and fourth pairs of pereiopods remained functional and moved in response to mechanical stimulation from a glass rod; however, given the forward pitch of the crab the viable limbs were unable to exert any force on the substratum and the crab could therefore be considered as functionally paralysed. Specific paralysis was not evident at concentrations below 20 mg L^-1^; however, differences in responsiveness relative to the controls were evident, even at the lowest assayed concentration (2 mg L^-1^). These non-paralysed crabs were noticeably more docile to the extent that the operator was able to manually remove them from the tanks at 72 hours and the crabs made little to no effort to resist beyond walking to the corner of their tank. Crabs exposed to sub-paralytic concentrations were therefore still affected and may be described as partially anaesthetised, but did not die within the 72 hour exposure.

### 4.2 Reactive oxygen species

There were no significant changes in cellular ROS (cROS) or lipid peroxidation ROS (LPO ROS) formation in the gills or claw muscle after 24 hours of exposure to low pyridine concentrations (<20 mg L^-1^; Figure 2A&C); however, only one of the three 20 mg L^-1^ crabs was alive after 24 hours, with the other crabs having died near to the 24-hour sampling time – this will likely have affected the overall ROS profile for that concentration. There was however a significant concentration dependent increase for mitochondrial ROS (mROS) in both the gills and claw muscle (Figure 2B). Despite concentration being a significant factor (Table 2), a Post-hoc Tukey test determined no significant differences between the controls and the three tested concentrations, likely due to the low number of replicates used.

**Figure 2.**
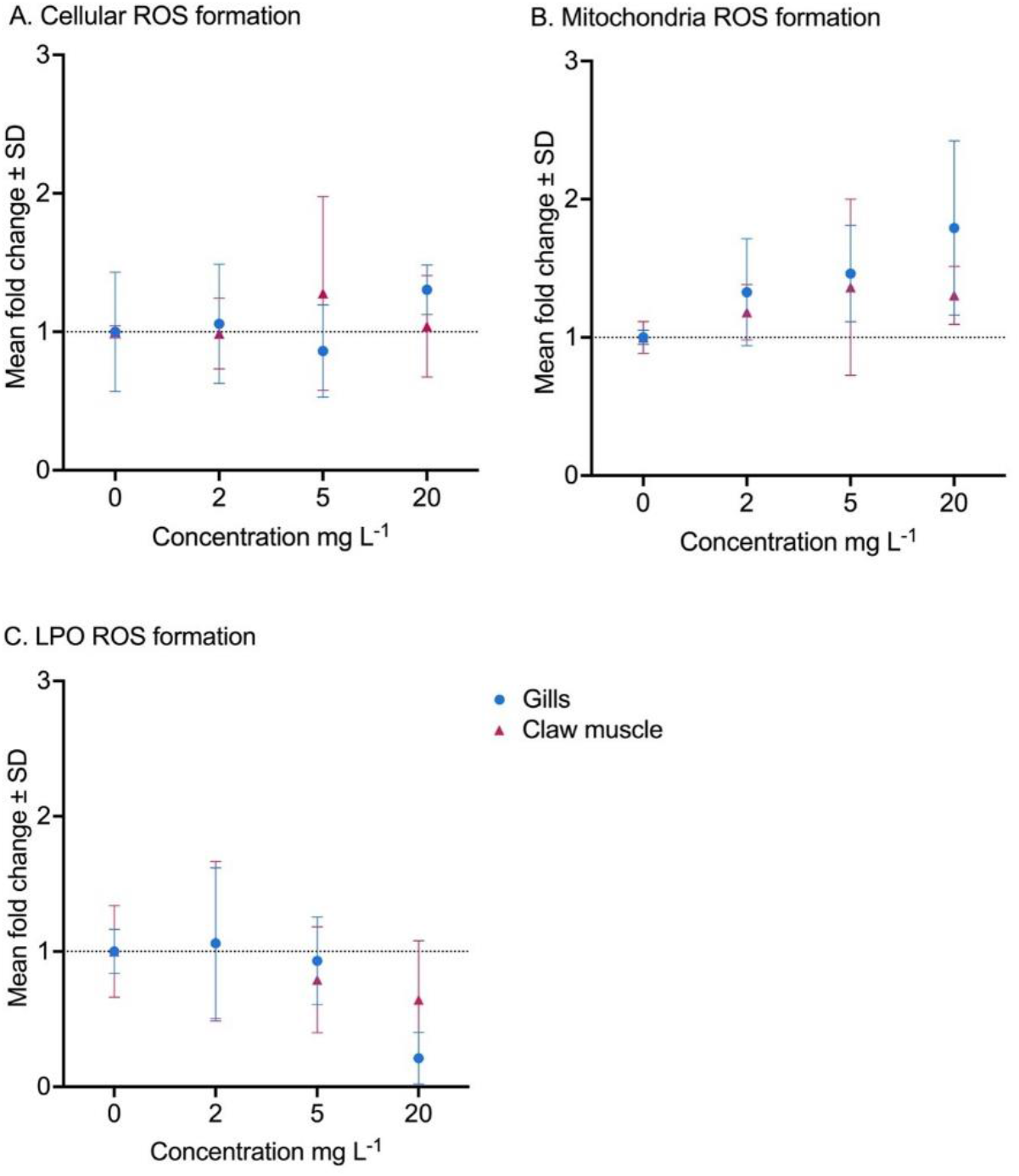
Formation of reactive oxygen species (ROS) in the (A) cells, (B) mitochondria and (C) lipid peroxidation (LPO) in *Cancer pagurus* after 24 hours of pyridine exposure. The blue circles represent the gills, and the purple triangles represent the claw muscle. The dotted line on the y-axis represents the control samples (fold change of 1).

Time was the only significant factor influencing cROS formation at 100 mg L^-1^ (Table 2), with a Post-hoc Tukey test revealing a significant difference between 3 and 6 hours for the claw muscle (Figure 3A). There was a considerable increase in mROS formation in claw muscle after 3 hours at 100 mg L^-1^, which returned to baseline levels at 6 hours (Figure 3B). As per the 20 mg L^-1^ treatment, the return to baseline levels for the ROS data will likely have been because all crabs had died by 6 hours at 100 mg L^-1^. Time, tissue and the interaction between time and tissue were all significant factors influencing mROS formation (Table 2). This pattern was also seen for LPO ROS, with all three factors being significant (Table 2) and a significant difference between 3 and 6 hours LPO ROS in the hepatopancreas (Figure 3C).

**Figure 3.**
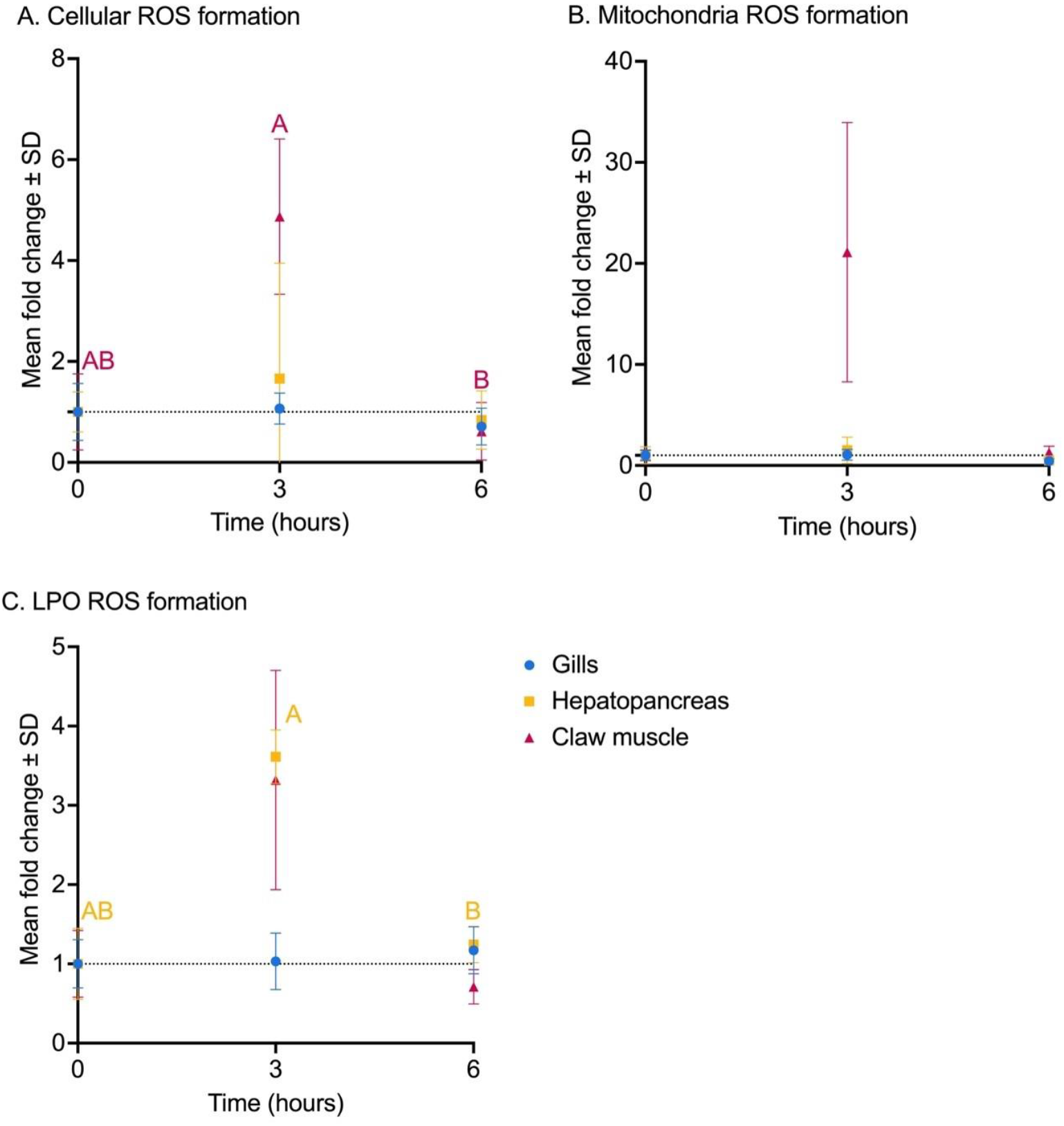
Reactive oxygen species (ROS) formation in (A) cells, (B) mitochondria and (C) lipid peroxidation (LPO) in *Cancer pagurus* after 3 and 6 hours of pyridine exposure at 100 mg L^-1^. The blue circles represent the gills, the yellow squares represent the hepatopancreas and the purple triangles represent the claw muscle. The dotted line on the y-axis represents the control samples (fold change of 1). The letters represent significant differences between time points and are colour-coded to the tissue they represent (P < 0.05).

### 4.3 Pyridine transport

Figure 4 displays the simulated total mass of pyridine in the domain against time (in kg, assuming a pyridine density of 982 kg m^-3^). Dredging and discharge operations stopped on the 3^rd^ of October 2021, at which point over 8000 kg of pyridine was predicted to be present in the water column (approximately half of the total amount released). An exponential decay ensues, with virtually no pyridine predicted to be left in the water column by about the 15^th^ of October 2021. Figures 5 and Figure S3 show the transport of the modelled pyridine plume predicted to have been released at both the dredging site and the spoil grounds. The plume is further represented as an animation in Figure S4.

**Figure 4.**
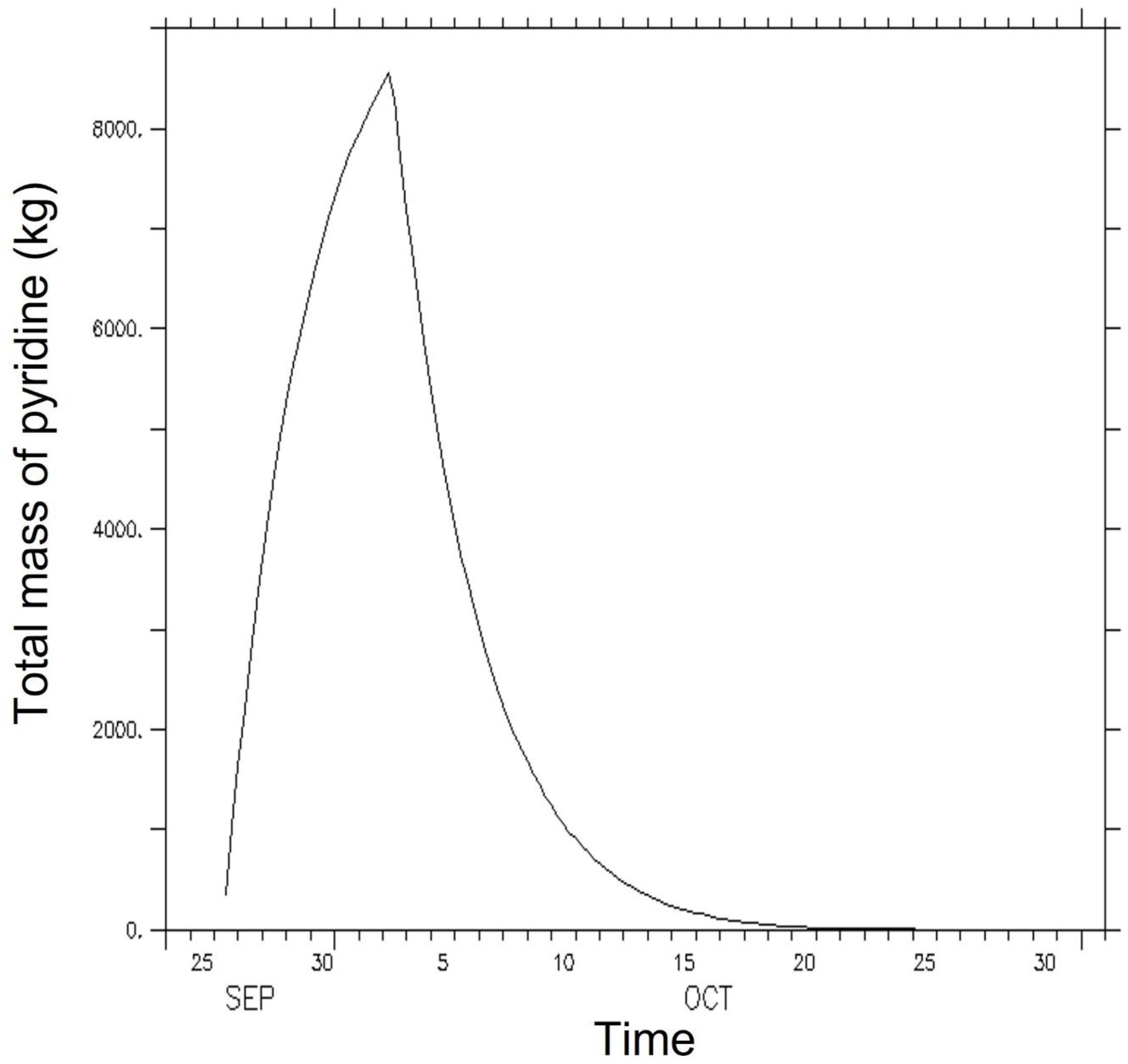
Total mass of pyridine in the entire model domain as a function of time, assuming a pyridine density of 982 kg m^-3^ and a half-life of 3.47 days.

**Figure 5.**
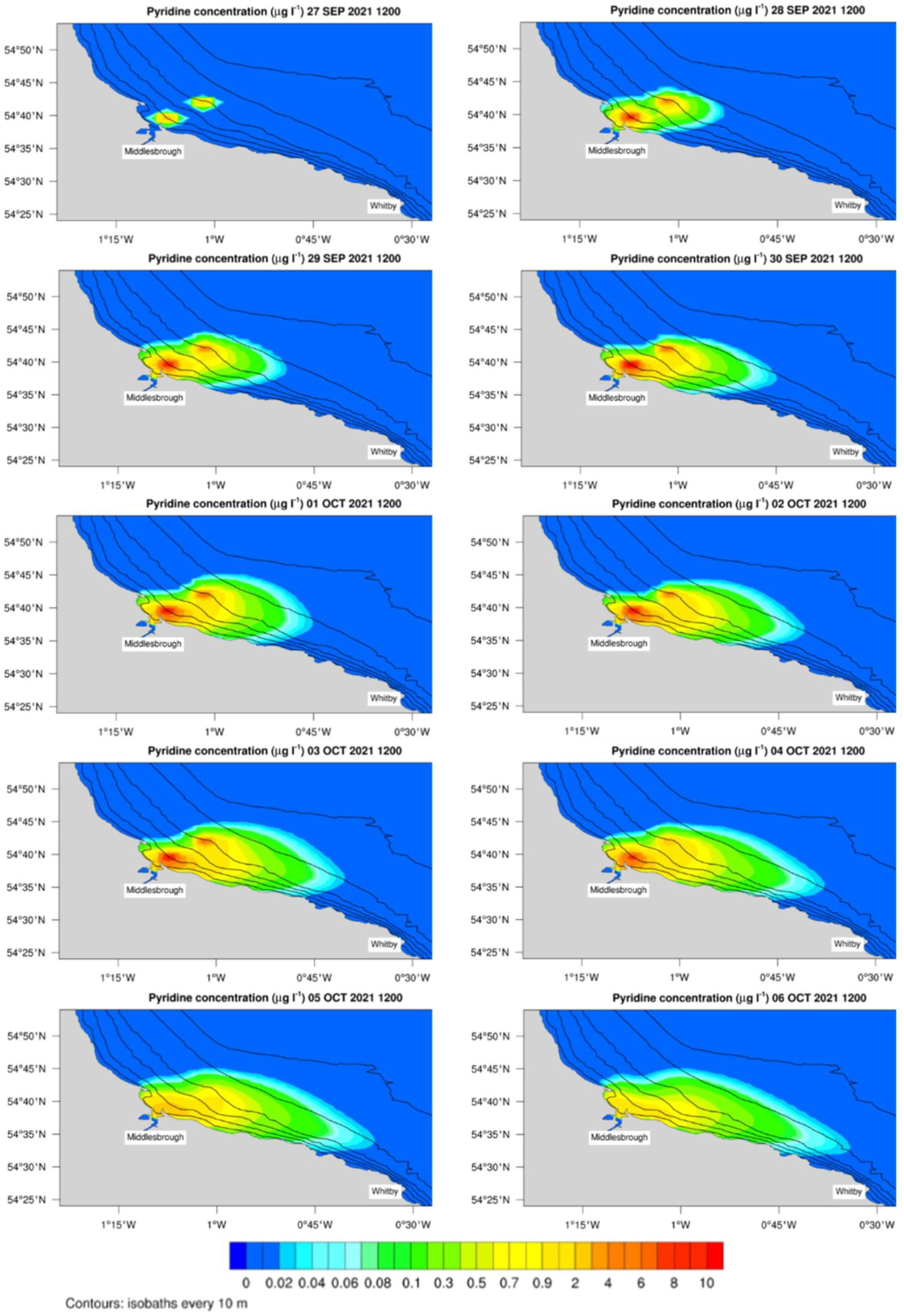
The predicted time course of the pyridine plumes released in the water column based on a simulated release of 10,000 L of pyridine at each of the dredge and spoil sites. These maximum concentrations are typically attained along or near the seabed. Pyridine decay is modelled with a half-life of 3.47 days, meaning that 90% of the released pyridine will have decayed after 8 days. The model predicted that pyridine would be transported to Whitby. This figure visualises the maximum pyridine concentration within the plume between the dates of September 27^th^ to October 6^th^, 2021. The position of the plume covering the dates of October 7^th^ to the 14^th^, 2021 are presented in Fig. S3.

Figure 6A shows the maximum predicted pyridine concentrations (μg L^-1^) in the water column, with the highest concentrations (>10 μg L^-1^) expected at the dredge site and readily dissipating as the plume is transported along the coastline. These maximum concentrations are typically attained along or near the seabed, but significant vertical mixing occurs as the plume spreads along the coast. While the dominant circulation leads to an overall dispersion of tracer towards the south east, tidal flows and eddy diffusion lead to mild propagation of tracer towards the northwest. We stress the important fact that the absolute concentration values shown in the figure are less meaningful than the temporal and spatial patterns displayed, because it is not possible at present to provide an accurate estimate of the possible amounts of pyridine that any dredging operations might have released into the water column. However, the dispersion of tracer is entirely plausible and agrees with our best current understanding of the circulation in the area as determined by the operational forecasts and analyses of AMM15. There is a clear potential over the period studied for plumes originating in the Teesmouth area to propagate as far south as Whitby.

**Figure 6.**
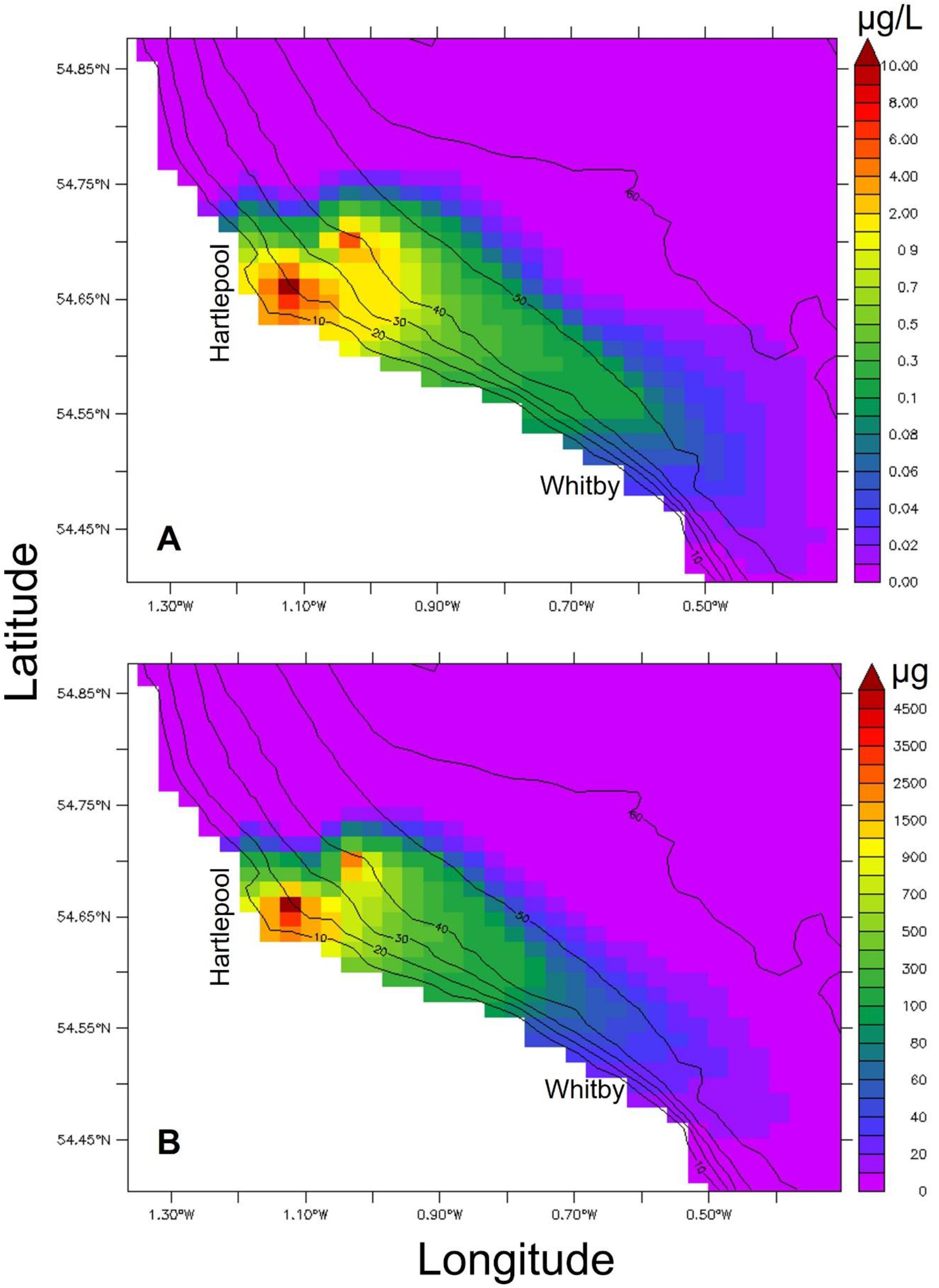
(Top) The maximum concentrations of pyridine (μg L^-1^) predicted to be typically attained along or near the seabed. (Bottom) The predicted cumulative exposure to pyridine (μg) for a nominal crab located throughout the pyridine plume.

Figure 6B shows the predicted cumulative exposure to pyridine for a given crab located at any point within the plume area. Cumulative exposure was calculated by postulating a volumetric ventilation rate (the volume of water flowing through the crab’s gill chamber) of 50 mL min^-1^ (O’Mahoney and Full, 1984). Note, this does not equate to bioaccumulation but does relate to the duration over which a crab would be exposed to pyridine, and therefore the period over which toxicity may be experienced. Highest cumulative exposure (>4500 μg) will have been experienced by crabs in the immediate vicinity of the dredge site, with crabs near the spoil dumping site predicted to have been exposed to up to 3000 μg. Cumulative exposure would reduce as the plume dissipates along the coast; however, the model predicts that crabs located off Whitby could still be cumulatively exposed to up to 50 μg of pyridine.

## 5 Discussion

Dredging of the River Tees has been suggested as a causative factor for mass die-offs of marine animals, notably among the decapod crustaceans (EFRA, 2022). Initial investigations suggest that pyridine (an industrial solvent with a long history of use and discharge into the Tees) may have played a major role, with exposure presenting as convulsions, paralysis, and death; however, key knowledge gaps remain. Chemically, pyridine has some potential for bioaccumulation. Its low octanol-water partition coefficient (log Kow = 0.57; (Cumming and Rücker, 2017) and low Henry’s law constant (log Kaw < −3) (Hawthorne et al., 1985), indicate that it may be mobile in water and is unlikely to rapidly volatilize into the atmosphere. As a weak base (Albert et al., 1948), it will be present predominantly in its neutral form in seawater, making it more likely to cross cell membranes. Pyridine is not oxidized rapidly in abiotic water (Mill et al., 1980) so its removal from an ecosystem is likely to depend on the rate at which it is biodegraded. A range of studies across different aerobic and anaerobic environments show that biodegradation half-lifes can be anywhere from a few days (Cassidy et al., 1988) to several months (Battersby and Wilson, 1989).

Moreover, pyridine is known to rapidly bioconcentrate during the first 1 to 2 hours of exposure in tissues of the guppy (*Poecilia reticulata*) before reaching an equilibrium; approximately 10% of compound uptake occurred during the first hour of exposure (de Voogt et al., 1991). If this uptake pattern holds true for other marine and aquatic organisms it would indicate the rapid build-up of a pyridine body burden that should be proportionate to the exposure concentration once equilibrium is reached (and assuming no depuration mechanism). Through very cautious use of this assumption, we can extrapolate this to the indicative measure of pyridine from the tissue of the dead crabs collected from the impacted sites around Teesmouth. The highest measured pyridine concentration was 439.6 mg/kg (using a non-validated method). Assuming a 10% uptake this would extrapolate to a localised pyridine exposure approximating to 4000 mg L^-1^. However, we urge caution in using this figure particularly as there remain unanswered questions surrounding the potential for post-mortem pyridine biosynthesis. Work to address this question is ongoing in our laboratory and as part of a wider research programme being undertaken by Cefas (EFRA, 2022).

There remains a paucity of toxicology data for aquatic organisms exposed to pyridine; examples include 73.6, 26 and 4.6 mg L^-1^ for 96-hour LC_50_ values for fathead minnow (*Pimephales promelas*), European carp (*Cyprinus carpio*), and rainbow trout (*Oncorhynchus mykiss*) respectively (ThermoFisher, 2009). The current work derived a 24-hour LC_50_ value of 19.44 mg L^-1^ and a 72-hour LC_50_ value of 2.75 mg L^-1^, which is an order of magnitude lower than for minnow and carp, but broadly comparable with trout. In unpublished work, we observed no evidence of toxicity to nauplii of the brine shrimp *Artemia* at concentrations up to 100 mg L^-1^ over a 72-hour exposure. In crabs, death was swift at lethal doses, i.e., within 6 hours of exposure at 100 mg L^-1^ with a consistent sequence of behavioural responses, progressing from convulsions and uncoordinated limb twitching into a paralysis stage, followed by death. At lower concentrations (<50 mg L^-1^) the convulsions were not observed, but twitching was still evident, albeit less pronounced. Behavioural change was noted at sublethal concentrations, even at the lowest concentration tested (2 mg L^-1^) wherein the crabs were noticeably more docile compared with control animals. It should be noted that the toxicity data reported here are likely to be underestimates. Pyridine is volatile and some loss from the exposure solutions would be expected. Given that the exposure tanks contained a significant headspace, and that volatilisation would likely have been accelerated through aerating each tank, it is reasonable to assume some loss of pyridine from our treatments. These issues were recognised and addressed by de Voogt et al. (1991) in their work and future exposure trials should seek to limit pyridine loss. Further, pyridine is degraded by oxidation and some degree of oxidative loss would be expected from our exposure trials (pyridine and pyridine derivatives are more stable and persistent in anaerobic sediment (Kuo and Liu, 1996;Liu et al., 1998)).

Survival and ROS formation are effect end-points that provide an indication of toxicity. ROS are cell signalling molecules (easily measured using fluorescent-probes, Gomes et al. (2005)) that are constantly produced within all organisms and converted into less reactive species. A build-up of ROS within a cell system can be a generalised indicator of localised stress, as systems are unable to keep up with detoxifying excess ROS (Paital and Chainy, 2012). Biochemical analysis revealed a complex picture relating to ROS production. There was no significant change in cellular ROS (cROS) at pyridine doses at or below 20 mg L^-1^ after 24 hours, however it should be noted that two of the three replicates exposed to 20 mg L^-1^ had died by 24 hours, therefore a loss of signal for that time point would be expected. Exposure to 100 mg L^-1^ triggered a significant spike of cROS production at 3-hours in the claw muscle but not in the gills or hepatopancreas. Interestingly, the mROS was highest in the gills at sublethal concentrations but there was no change at 100 mg L^-1^; rather, there was an increase in mROS production in the claw muscle. This observation corresponds with the widespread autotomy of claws and legs in crabs exposed to 100 mg L^-1^ (this was also commonly observed at 50 mg L^-1^ although ROS data were not collected for this concentration). There was no significant change in lipid peroxidation ROS (LPO ROS) production at concentrations at or below 20 mg L^-1^, however there were significant increases in both the claw muscle and hepatopancreas at 3 hours when exposed to 100 mg L^-1^.

The data demonstrate that pyridine is highly toxic to *C. pagurus*, ranging from distinctive patterns of acute toxicity to observable behavioural responses at much lower concentrations. The observed behaviours under acute exposure correspond with reports from the fishing community and other coastal users during the mass mortalities, specifically that moribund animals were presenting as twitching and paralysed. Further, animals that survived the initial impact of the autumn 2021 event continued to die over a longer period, resulting in a sequence of progressively smaller wash-ups and widespread reports of crabs and lobsters dying in holding tanks used by various actors along the seafood supply chain.

We extrapolate our findings in crabs to the other decapod species impacted by the 2021 autumn mass mortalities. Our LC_50_ data indicates that *C. pagurus* is more susceptible to pyridine toxicity than other commonly used bioassay organisms (fish and non-decapod crustaceans). Pyridine is structurally related to a family of toxic pyridine alkaloids synthesised by nemertean worms; for example, anabaseine and 2,3’-bipyridyl (the latter is also known as the crustacean convulsant factor) (Kem et al., 1976;Kem, 1985;Kem and Soti, 2001;Göransson et al., 2019). These pyridine derivatives are ecologically bi-directional in that they are released into the worm’s mucus to deter predation by crustaceans (thus functioning as a toxungen or crinotoxin), and simultaneously form part of the worm’s venom that is used to facilitate predation on crustaceans and polychaete worms. The pyridine alkaloids are reported to target the nicotinic cholinergic receptors in the crustacean central nervous system and the stomatogastric muscle nicotinic receptor calcium channel (Göransson et al., 2019). It has been reported that spiny lobster (*Amphiporus angulatus*) will rapidly withdraw upon contact with nemertean mucus and subsequent feeding is only observed if the mucus is cleaned from the worm (Göransson et al., 2019). This response strongly implies some form of external and rapid detection of the pyridine compounds. Interestingly, the evolution of pyridine analogues as kairomones is gaining increasing attention well beyond the nemertean-crustacean model (Brechbühl et al., 2015).

Crayfish are known to possess pyridine-sensitive neurons (also referred to as pyridine-sensitive units or pyridyl-sensitive neurones), primarily located on the pereiopods. These neurones are highly sensitive to pyridine and pyridine derivatives (Altner et al., 1983;Hatt and Schmiedel-Jakob, 1984;Carr et al., 1987), and show some degree of temperature dependence in terms of action potential frequency but not in terms of the spontaneous initiation of activity (Hatt, 1983). Hatt and Schmiedel-Jakob (1984) produced a sensitivity matrix as part of their comprehensive survey of substances stimulatory to pyridine-sensitive neurones in the walking legs of the crayfish *Austropotamobius torrentium*. The most stimulatory compound was pyrazinecarboxamide, with a K_m_ of 1.5 x 10^-6^ mol L^-1^. Pyridine was determined as the 12^th^ most stimulatory compound with a K_m_ of 4 x 10^-4^ mol L^-1^ – this equates to an exposure concentration of 31.64 mg L^-1^, i.e., comfortably within the toxicity range identified in the current study. This indicates that pyridine, above some threshold concentration, is functioning as a neurotoxin. 4-aminopyridine, which Hatt and Schmiedel-Jakob (1984) identified as an order of magnitude less stimulatory than pyridine, is known to rapidly block the potassium currents in the giant axon of *Carcinus maenas* and *Cancer pagurus* when applied either internally or externally (Quinta-Ferreira et al., 1982;Soria et al., 1985).

Pyridine and pyridine derivatives play major roles in the chemical signalling and environmental responsiveness of decapod crustaceans, with clear scope for neurotoxicity if safe environmental threshold concentrations are exceeded. Despite the structural relatedness with the pyridine alkaloids, the precise target(s) of pyridine in the crustacean central nervous system remain(s) undefined. While there is little proteome information on the affected UK crustacean species, in the American lobster (*Homarus americanus*) the Cys-loop receptors with immune-related domains may represent a link between immunity and neural signalling (Polinski et al., 2021), with some of the novel receptors highly expressed in the brain. Much work remains to be done to precisely define the range and location of pyridine receptors in these crustaceans. Further, there may be scope for pyridine to play a role in secondary toxicity by forming adducts with nicotinamide adenine dinucleotide (NAD^+^), which is a coenzyme pivotal to cell metabolism, and inhibiting enzymes requiring NAD^+^, involved in pathways key for survival. Adducts of NAD^+^ and small heteroaromatic molecules such as pyrinuron, pyridine, and 3-acetylpyridine, formed through base-exchange reactions (Fig. S5), have been recently reported as potent inhibitors of mammalian sterile alpha and toll/interleukin-1 receptor motif-containing protein 1 (SARM1), which cleaves NAD^+^ (Sarkar et al., 2021;Wu et al., 2021;Shi et al., 2022). Human SARM1 can catalyse NAD^+^ adduct formation with pyridine and multiple pyridine-fused heterocycles and nicotinamide mimetics either as purified protein or in neurons (Shi et al., 2022). A similar base-exchange mechanism has been described earlier for the human NAD^+^-ase CD38 (Preugschat et al., 2008). Recently, structurally- and functionally-related proteins have been reported in crustaceans, including toll-like receptors (TLRs) and SARM homologues (Kim et al., 2017;Habib and Zhang, 2020). These have been described to exert important roles in host defense of crustaceans against their pathogens, but it is likely that these proteins exert other roles, much like their mammalian counterparts, that remain undiscovered. Our currently ongoing investigation aims to determine whether NAD^+^-pyridine adducts may indeed be detected in crabs, and to elucidate plausible molecular targets of pyridine responsible for the observed toxic effects.

Given the clear toxicity of pyridine to crabs, we applied computational approaches to simulate the rate and extent of pyridine transport on the assumption that it was released from anoxic sediment as part of the UKD Orca dredging campaign. As part of the DEFRA joint agency investigation into the mass mortalities, Cefas undertook some initial modelling of the transport of pyridine from the spoil dumping site (DEFRA, 2022). These models only factored in the spoil site and not the dredging site. Further, the models were only run for a 3-day simulation. We decided to extend Cefas’ initial modelling work by factoring in pyridine release from the dredge and spoil sites and by running our model for a 30-day period. The simulated release of pyridine (10,000 L) was the same across both models; albeit that the Cefas model only released pyridine at the spoil site. Further, our model incorporated a pyridine removal function based on an assumed loss of 90% over an 8-day period (ATSDR, 2011). The model demonstrated that pyridine would have predominantly been transported southwards and would have reached Whitby and Robin Hood’s Bay in North Yorkshire. There would also have been more northerly transport (primarily due to tidal action), with potentially toxic levels predicted to reach Seaham in Co. Durham. The predicted transport of pyridine corresponds very well with the geographical range of the wash-ups, and with a gradation of impacts, with severe impacts felt around Hartlepool, Teesmouth and Redcar, and progressively reducing as the plume was transported southwards. The cumulative exposure model predicted that crabs off Hartlepool, Teesmouth and Redcar could have been exposed to a cumulative pyridine dose of >4.5 mg, with crabs sited mid plume, e.g., at Staithes and Runswick Bay being exposed to 300-500 μg, and crabs off Whitby being exposed to up to 50 μg of pyridine. These values must, for the time being, be treated cautiously as the exact volumes of pyridine assumed to have been released from the dredging work remain unknown.

Despite the strong fit of our model to on the ground reports of wash-ups, there remain areas in which our model needs refinement. While release of the pyridine tracer near the seafloor is probably a realistic representation of events when the vessel is discharging its load, it is possible that sediments and debris are injected more uniformly through the water column while dredging is occurring. Vertical distributions of tracer release should therefore be simulated. Barring the use of a more sophisticated model for the chemistry of pyridine sorbed to the sediments and dissolved in water, different pyridine decay rates should be investigated, notably since pyridine sorbed to solid particles will be generated both upon dredging and discharge may have a longer half-life than dissolved pyridine. Volatilisation of dissolved pyridine at the air-sea interface and deposition of pyridine back onto the seafloor as the sediment plumes settle should also be considered.

## Supporting information

Fig S4 animation

## Acknowledgements

We thank the crew of the RV Princess Royal for help with sourcing crabs; and Tony Baker and Peter McParlin for technical support. We are grateful for funding from The Fishmongers’ Company’s Fisheries Charitable Trust: (Charity number 284888).

## Ethics

This work complies with the ARRIVE guidelines for the use of animals in research and was carried out in accordance with the UK Animals (Scientific Procedures) Act, 1986 and EU Directive 2010/63/EU for animal experiments. This study was approved by the Newcastle University Ethics in Research Committee.

## Declaration of competing interests

JR is a founding member of the North East Fishing Collective which is an organisation representing the commercial fishing associations, angling societies, and stakeholders along England’s north east coast that have been impacted by the mass mortality events. The remaining authors declare no competing interests.

