## Supplementary figures for "Determining the toxicity and potential for environmental transport of pyridine using the brown crab *Cancer pagurus* (L.)"

*Corresponding author

**Supplementary Materials**

**Figure legends**

**Figure S1.** The NEMO (Nucleus for European Modelling of the Ocean) model domain covering the spherical rectangle (48^o^N,3^o^W)–(58^o^N,3^o^W). The integration grid is identical to that of the Atlantic Margin Model 1.5 km, AMM15 for that area.

**Figure S2.** (Left) A representative crab exposed to a pyridine concentration of 100 mg/L. At this concentration the crab presented convulsions before the onset of paralysis. (Right) The only surviving crab exposed to a pyridine concentration of 20 mg/L after 72 hours. Despite being alive that crab presented partial paralysis.

**Figure S3.** The predicted transport of pyridine released during the September and October 2021 dredging campaign. This figure visualises the plume between the dates of October 7^th^ to the 14^th^, 2021.

**Figure S4.** Animation of the release and transport of pyridine from dredge and spoil dumping sites.

**Figure S5.** Proposed mechanism for pyridine adduct formation. Pyridine (PYD) undergoes base-exchange reaction with nicotinamide adenine dinucleotide (NAD^+^), yielding an adduct and free nicotinamide (NAM).

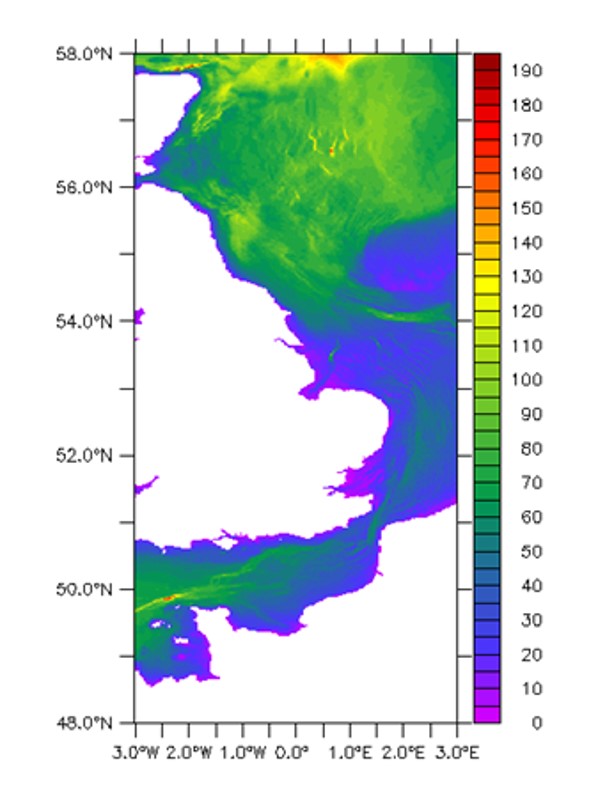

Fig. S1.

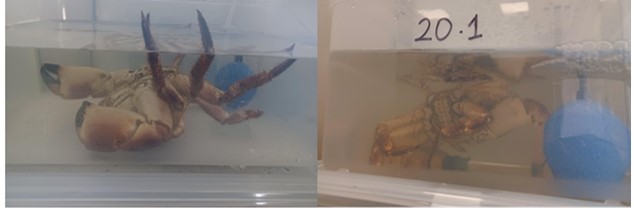

Fig. S2.

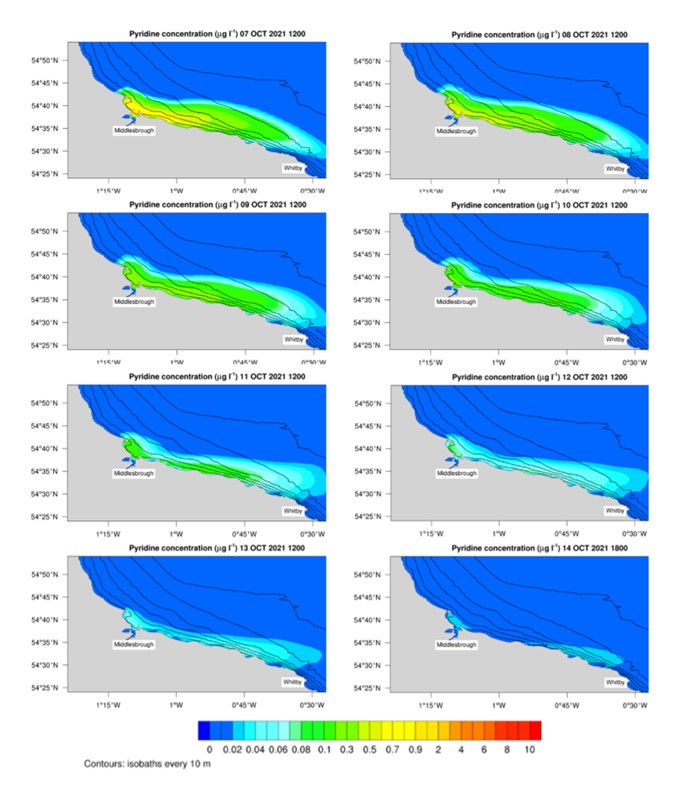

Fig. S3.

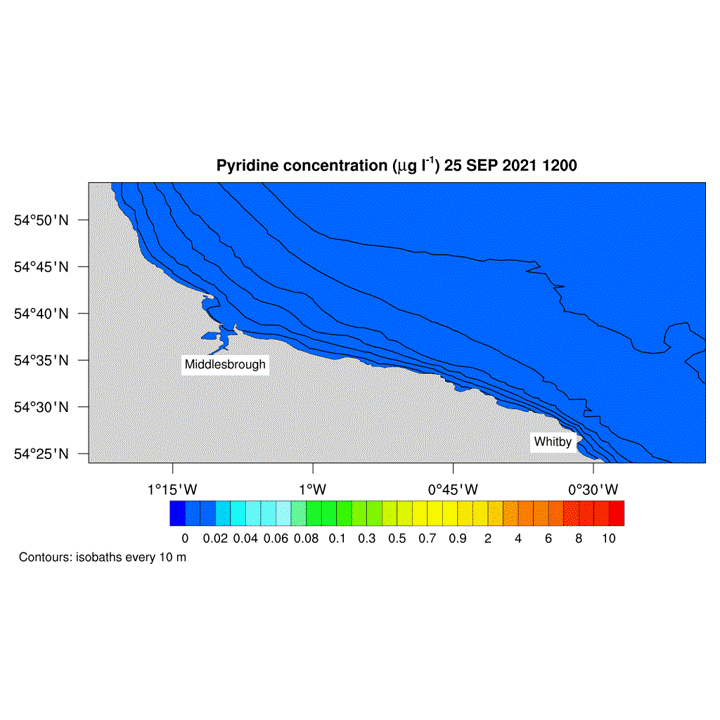

Fig. S4.

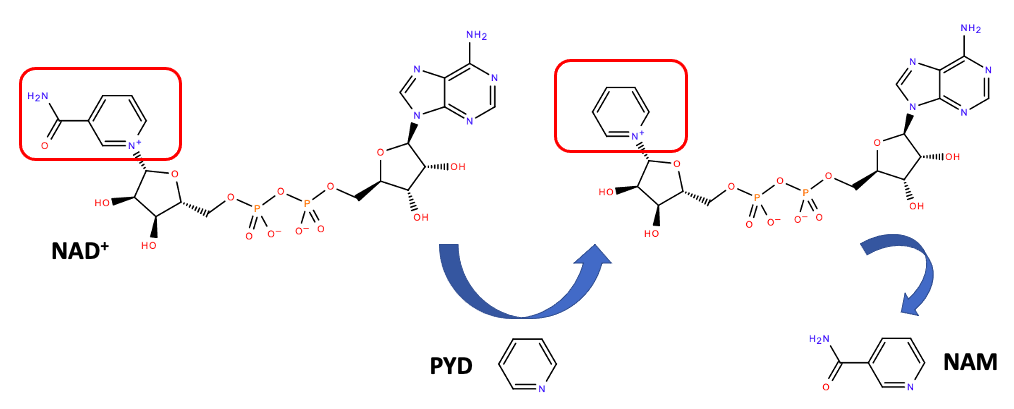

Fig. S5.
